# Simple rules for an efficient use of Geographic Information Systems in molecular ecology

**DOI:** 10.1101/113225

**Authors:** Kevin Leempoel, Solange Duruz, Estelle Rochat, Ivo Widmer, Pablo Orozco-terWengel, Stéphane Joost

**Affiliations:** Ecole Polytechnique Federale de Lausanne (EPFL), School of Architecture, Civil and Environmental Engineering (ENAC), Laboratory of Geographic Information Systems (LASIG), CH 1015 Lausanne, Switzerland; Cardiff University, Biomedical Science Building, Museum Avenue, Cardiff CF10 3AX, Wales UK

**Keywords:** Geographic Information Systems, Spatial Analysis, Landscape Genetics, gene-environment associations, Open-source software, Geographic map

## Abstract

Geographic Information Systems (GIS) are becoming increasingly popular in the context of molecular ecology and conservation biology thanks to their display options efficiency, flexibility and management of geodata. Indeed, spatial data for wildlife and livestock species is becoming a trend with many researchers publishing genomic data that is specifically suitable for landscape studies. GIS uniquely reveal the possibility to overlay genetic information with environmental data and, as such, allow us to locate and analyze genetic boundaries of various plant and animal species or to study gene-environment associations (GEA). This means that, using GIS, we can potentially identify the genetic bases of species adaptation to particular geographic conditions or to climate change. However, many biologists are not familiar with the use of GIS and underlying concepts and thus experience difficulties in finding relevant information and instructions on how to use them. In this paper, we illustrate the power of free and open source GIS approaches and provide essential information for their successful application in molecular ecology. First, we introduce key concepts related to GIS than are too often overlooked in the literature, for example coordinate systems, GPS accuracy and scale. We then provide an overview of the most employed open-source GIS-related software, file formats and refer to major environmental databases. We also reconsider sampling strategies as high costs of Next Generation Sequencing (NGS) data currently diminish the number of samples that can be sequenced per location. Thereafter, we detail methods of data exploration and spatial statistics suited for the analysis of large genetic datasets. Finally, we provide suggestions to properly edit maps and to make them as comprehensive as possible, either manually or trough programming languages.

## Introduction

Geographic Information Systems (GIS) are powerful tools to be used in the context of evolutionary studies. They are designed to store, handle, display and analyze any kind of data representing objects (individuals, populations, areas, etc.) characterized by geographic coordinates (X = longitude and Y = latitude). With the help of GIS, geographic information can be combined with for example phenotype, genotype, or environmental data to display the spatial distribution of genetic variants and to visualize factors influencing spatial evolutionary processes. The main advantage of GIS in evolutionary biology is to easily explore and display genetic information (neutral and adaptive genetic variation, gene flow) at multiple scales, and to overlay this information with physical barriers, land cover or topographic maps in order to generate subsequent analyses regarding the location and causes of genetic boundaries (Epperson, 2003; Manel and Holderegger, 2013). GIS have also proven useful in adaptive landscape genomics or gene-environment associations (GEA) studies, in the context of which they enable the retrieval of environmental variables at sampling locations. The integration of GIS with approaches from landscape ecology and population genetics, defined as landscape genetics by Manel et al. (2003), also has important implications for conservation biology (Petren, 2013).

However, since a few years we face a paradigm change with the advent of Next Generation Sequencing (NGS) data, whose use requires rethinking study pipelines. First, NGS currently presents an economic constraint as it is costly compared with genetic markers produced so far (e.g. microsatellites and AFLP). Consequently, we are unable to fully sequence several hundreds of individuals, requiring a careful selection of the samples to analyze. Appropriate sampling is thus key to achieve a precise and continuous evaluation of environment-driven selection on the genome (Hand et al., 2015; Manel et al., 2012; Rellstab et al., 2015). In addition, NGS datasets are large and must be treated differently to be efficient and to avoid computer memory overload. Finally, NGS data require new tools to display and analyze spatial patterns that are more computationally demanding.

The successful application of GIS tools is not intuitive for many biologists who are not familiar with the concepts relating geographic information systems and the use of GIS software. Indeed, a large diversity of GIS tools is available and the difficulty of finding relevant information and instructions is an obstacle for non-expert users. To date few scientific articles have defined the role of GIS in molecular ecology. For instance, Kozak et al., (2008) review the fast development of GIS-based environmental data and advocate for their usage as an alternative to unprecise proxies such as latitude of distance between populations. Another review by Joost et al. (2010) provided guidelines for GIS use in livestock genetics and enumerate the advantages of integrating data in a GIS environment. More recently, Rogers and Staub (2013) outline spatial analyses and GIS methods in honey bees research. Their review is not specific to bees but instead aim to intensify the exploration of the spatial component of studies in ecology and related disciplines. Lastly, Balkenhol et al. (2015) published a book detailing the concepts and analytical steps of landscape genetics studies, such as sampling design, spatial analysis and environmental datasets. Also, GIS are exploited in many unrelated domains and it is thus difficult to find resources specifically targeted at biologists. The bases of GIS practices are readily found in freely available Massive Open Online Courses (MOOCs), such as the Coursera platform (Coursera, 2012) currently offering six courses on GIS. Yet, these reviews do not tackle the challenges brought by large genetic and environmental datasets, and fail to review the recurrent caveats related to spatial research. In this paper, we highlight the usefulness of GIS in population and landscape genomics and provide key information for their successful application to these fields.

## Geographic coordinates

Geographic coordinates of samples constitute an invaluable source of information, ranging from the display of their spatial distribution to the retrieval of environmental variables. Whenever doing fieldwork, using a GPS is the best way to record the coordinates of samples. As such, we strongly recommend recording the location of each sample, instead of the location of the centroid of a population for instance. Firstly, it allows for a more precise retrieval of environmental values. Secondly, attributing the same location to several samples invokes pseudo-replication, a statistical bias that must be addressed in further analysis. Thirdly, coordinates of nearby individuals allow for a proper measurement of dispersal, using for example pairwise genetic relationship with distance. Regarding GPS devices, standard GPS, and to a lesser extent smartphones, are accurate enough in most cases. However, more precise devices, such as DGPS (differential GPS), are recommended for local scale studies in which samples are located less than a couple of meters apart: the precision of the location has to stay within the spatial resolution of the grain.

When GPS coordinates are not recorded, it is still possible to approximate sample locations with the help of satellite images or by encoding the address of the location (georeferencing or geocoding), although with a lower accuracy. In the former case, creating a new vector layer overlaid on a satellite image or an online map (see next section) allows the recovering of samples coordinates from an approximately known location (e.g. a crossroad, a tree, a river; Docs.QGIS, 2014). For the latter case, plugins have been developed to read text delimited file containing addresses (e.g. your own house address) that you want to locate (for example the MMQGIS plugin in QGIS, Mangomap, 2012, MMQGIS plugin, 2012)). It must be noted that each line must contain the address, city, state and country.

Another essential consideration is choosing the relevant coordinate reference system. Indeed, GPS devices display the coordinates of a point in latitude and longitude values, usually in the World Geodetic System (WGS84). This is a global reference system in which the Earth is represented by an ellipsoid, and every position on the surface is defined by two angles at the center of the Earth: the latitude and longitude. However, projected systems for which a geographical location is converted from the ellipsoid (distances expressed in degrees) to a corresponding location on a two-dimensional surface (x and y expressed in meters) are preferred for analyses. It is important to note that, although global systems covering the whole planet exist, each country or region has its own coordinate system that is locally more accurate than the global system. Where no national projected system exists, it is still possible to use the Universal Transverse Mercator (UTM) coordinate system, a projected coordinate system covering the entire globe and dividing it into sixty 6°-wide longitudinal zones (Dmap, 1993). Even though GIS software usually deal with different projection systems, the manual reprojection of all layers into the same local projection system is recommended to avoid potential incompatibilities (see next section). However, different GIS may not exactly use the same name for a coordinate system. Therefore, to facilitate the identification of coordinate systems across the diversity of GIS software, the EPSG (European Petroleum Survey Group) database (EPSG, 1985) is a widely used database referencing all projected coordinate systems, implemented in every GIS and providing them with a unique ID (Maling, 1992), e.g. EPSG: 4326 correspond to the WGS84 reference system.

## Software

There are many GIS software, with different functions and aimed at various audiences. Universal GIS software do not exist and, therefore, the choice is difficult for a beginner. Today, one of the most user-friendly GIS is QGIS (QGIS Development Team, 2015). It is ideal to explore geodata, able to read and convert a wide variety of input formats and suitable to produce high-quality maps. Note that QGIS, and all other GIS mentioned in this paper, is free and open source. Open source GIS can indeed perform the same tasks as their commercial counterparts, and include the opportunity to understand and improve GIS algorithms or enable a better collaboration as there is no problem related to license access (Ertz et al., 2014). In addition, a large community exists to support development efforts of open source GIS, and regularly creates extensions to add functions and improvements to the software. Forums and tutorial websites are also flourishing for newcomers (http://gis.stackexchange.com/, http://www.qgistutorials.com/, http://www.qgistutorials.com/, Sutton et al., 2009).

On the other hand, most analysis in GIS are not easily replicable and, therefore, programming languages such as R can be more efficient. R has been successfully used as a GIS for a long time and several packages and reviews have been published (Brunsdon and Comber, 2015; Rodriguez-Sanchez, 2013).

Among them, we can mention *rgdal* for the importation of geodata (Bivand et al., 2016), *GISTools* for general GIS operations (Brunsdon and Chen, 2014), *rasters* for their display (Hijmans and van Etten, 2015), *spdep* and *spatstat* for spatial statistics and analysis (Baddeley and Turner, 2005; Bivand and Piras, 2015). While these packages are relatively efficient to import, display large rasters and vectors, customization options are more limited than in dedicated GIS.

## Main dataset

The first step in a GIS project is usually to import a vector file containing samples coordinates. QGIS has a plugin to easily import GPS coordinates, either directly from a GPS device or through vector files, such as .kml or .gpx. (Docs.QGIS, 2013). These formats are usually converted to shapefiles (.shp) due to the easier management of their attributes and projection system associated with vector units. Delimited text files (e.g. tabulator - tab - or space delimited) can be easily opened in QGIS as well and then be transformed into shapefiles. When opening a text file using “Add delimited text layer”, QGIS should recognize automatically the delimiter used and the columns of coordinates (X Y, Latitude Longitude) (QGIStutorials, 2014). However, such delimited text files cannot be transformed to polygons or lines. In this case, one should already have a shapefile incorporating lines or polygons to which the text table can be joined. To do so, the shapefile and the text table should have the same column of unique IDs. When clicking on the properties of the shapefile, an option is proposed to join additional tables of attributes (QGIStutorial, 2014). As mentioned in the previous section, it is recommended to project all layers in the same coordinate system. In QGIS, this is done by right-clicking on the layer and by changing the coordinate system in the “save as” option. The newly projected layer will then be automatically loaded to the project. See Rogers and Staub (2013) for a more extensive review of the basic tasks in QGIS.

## Background dataset

The second step is to add one or more background layer(s) to constitute the geographic context, either from raster data (see next section) or from an online map (Google, Bing, Open Street Map). The OpenLayer plugin in QGIS allows the addition of a background base-map to the QGIS interface (QGIS workshop, 2013). When using raster layers such as Elevation data or climatic variables, adding a semitransparent shaded relief will enhance the contrast and reveal the topography. To this end, QGIS has a Terrain analysis module in which a hill-shade layer can be computed from a Digital Elevation Model (DEM, i.e. a matrix of elevation data). Then, the transparency of the layer can be adjusted in its properties. In addition, it is advisable to cut rasters and vectors to the size of the study area using the clipper tool to facilitate their display and reduce computation time. Note that the succession of layers in the main frame depends on the order of layers shown on the left panel of the application.

## Environmental and landscape variables

Environmental datasets have considerably evolved and represent new opportunities for the identification of environmental drivers of adaptation. One of the main applications of GIS software in landscape genomics is to extract values of environmental variables at the exact location where samples have been collected, or from the surrounding area by means of polygons representing a buffer, a forest, or a specific land cover class for instance. As databases containing georeferenced environmental variables are numerous, we propose a list of the 10 most important publicly accessible databases in Table 1 (A more extensive list is proposed in Appendix 1). Raster environmental data are often delivered in geotiff (.tif) or Band Interleaved by Line (.bil) formats, similar to satellite images but containing only one layer of information (i.e. Temperature, Precipitation etc.). Regarding climate datasets, many studies rely on variables interpolated at large geographical scales on the basis of data provided by weather stations and distributed across territories, such as the WorldClim dataset (Hijmans et al., 2005). These data are often delivered as continuous grids and their spatial resolution (i.e. area covered by a pixel) typically varies between 1 and 10 km2. For more local or regional databases, however, national agencies are the most valuable sources.

**Table 1:**
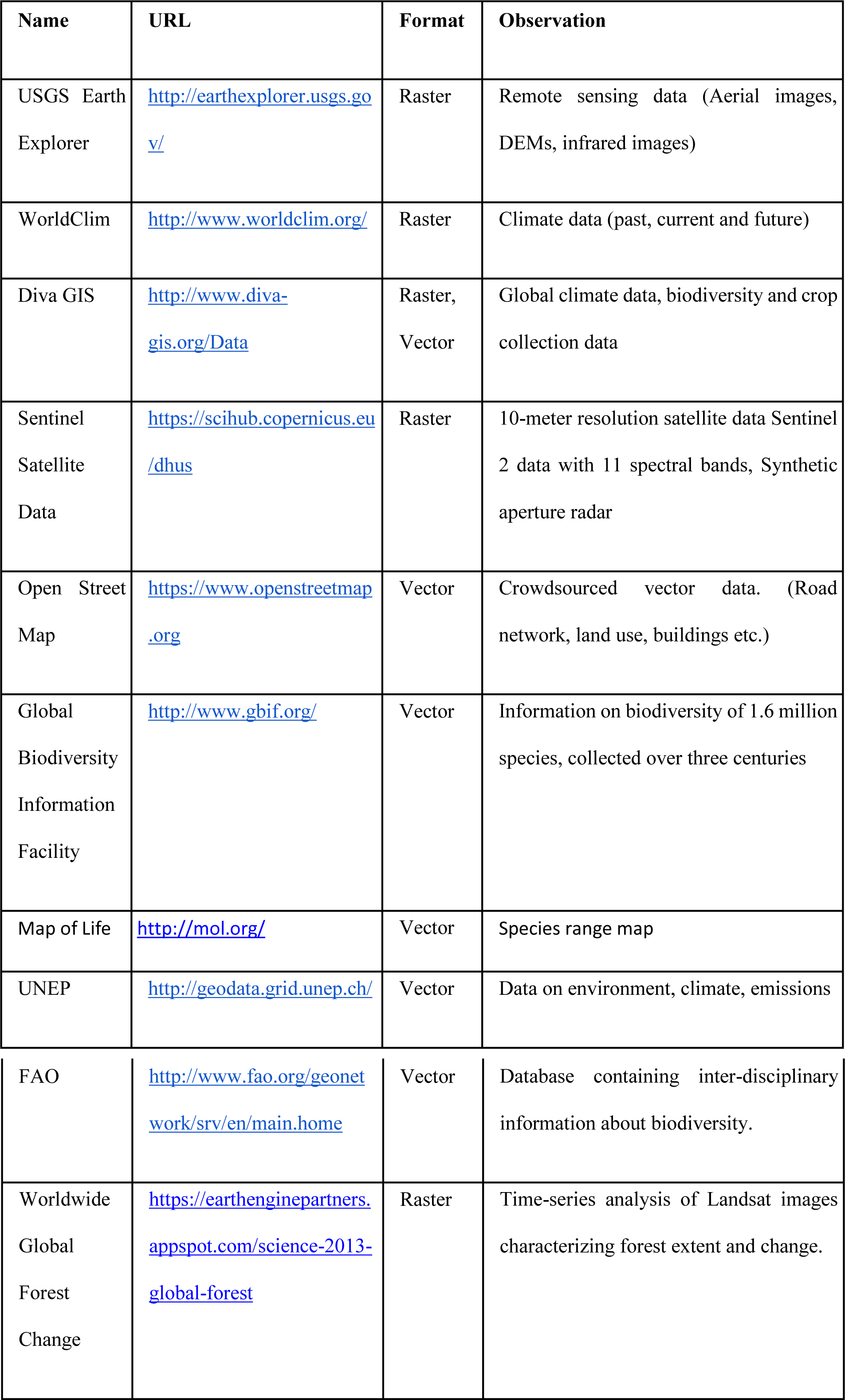
10 main public sources of environmental data (URLs: consulted on June 10, 2016)

Alternatively or additionally, environmental variables can be computed from DEMs, and used as proxies to relevant ecological features (Kozak et al., 2008; Leempoel et al., 2015; Manel et al., 2010). DEMs are available on Earth Explorer (Earth Explorer, 2016) and come in formats such as geotiff or SAGA Grids (.sgrd). We recommend not using text formats for grids (such as .asc or .xyz) since DEMs resolution has dramatically increased over the years, making these formats slow and heavy. The most common use of DEMs in ecology consists in retrieving altitude or computing primary terrain attributes (i.e. slope, orientation and curvature) (Guisan and Zimmermann, 2000; Kozak et al., 2008; Manel et al., 2010; Wilson and Gallant, 2000). However, we recommend going beyond the traditional use of DEMs as a diversity of variables can be computed, like e.g. solar radiation, morphometric indices or hydrological variables (Leempoel et al., 2015). The treatment of DEMs and the production of topographic variables can be processed in software like SAGA GIS (Böhner et al., 2006; SAGA GIS, 2004) or GRASS GIS, now included in QGIS. SAGA GIS is the most DEM-oriented GIS to date and can compute a large panel of derived variables. It is also easily scriptable both using the command line or the R package RSAGA (Brenning, 2008), although the former is faster.

Satellite images covering the whole surface of the globe are also available through Earth Explorer and can be used e.g. to produce land cover maps. Most satellite sensors provide images with more than the 3 visible “colors”, or bands, and it is thus the choice of the user to decide which bands to attribute to color channels (Red, Green and Blue). For example, by assigning the infrared band to the red channel and green and blue bands to their respective channels, one can easily identify trees or forests against water, fields or naked soils because plants reflect infrared wavelengths more than other land cover types. This process, the supervised classification of remote sensing data (satellite and aerial images, radar, etc.), can be operated in Opticks (Opticks, 2001) or in SAGA GIS.

Finally, vector databases, such as Openstreetmap (OSM) (OpenStreetMap, 2004), are interesting in order to recover road networks, rivers, watershed boundaries, or landuse. OSM data can also be easily accessed through GeoFabrik (GeoFabrik, 2011) where cities or countries are already extracted. Note however that OSM data and most tiled web-maps are provided in Pseudo-Mercator projection (EPSG: 3857).

It is worth mentioning that in GEA studies, using a wide range of environmental variables often implies redundancy between these variables. However, statistical analyses require independent variables and, for this reason, it is important to perform multi-collinearity analysis (e.g.) on the set of environmental variables, to understand which variables are highly correlated (Dobson and Barnett, 2008; Fischer et al., 2013). Such collinearity can be detected by performing a PCA, by using Variance Inflation Factor (VIF) or calculating pairwise correlation coefficients between pairs of variables, and then removing randomly one of the two variables from a pair that shows high correlation. See Rellstab et al., (2015) for a review of these methods. However, bear in mind that environmental variables, in particular DEM-derived ones, may not have a normal distribution. Variables should thus be transformed or non—parametric tests should be used to test for correlations (for example Spearman ranks instead of Pearson correlation coefficients).

## Spatial analysis

Numerous spatial analysis techniques have been developed to address issues related to spatial data (Fortin and Dale, 2005). Here we focus on exploratory spatial data analysis (ESDA) and spatial statistics given their central role in molecular ecology. For other spatial analysis methods, we suggest to have a look at the Geospatial analysis guide (Smith et al., 2005) and at the spatial analysis guide for ecologists (Fortin and Dale, 2005).

Evolutionary biology can benefit from ESDA (Joost, 2006), an interactive approach allowing the user to explore and analyze a dataset dynamically and in real-time through a combination of various tools for data representation (Anselin, 1994). For example, maps can be used to display the position of samples, histograms and boxplots to evaluate the distribution of attribute values and Moran's scatter plot or conditional plots to analyze the relationship between the various variables. ESDA can also be useful for example to localize samples in areas showing extreme climatic conditions (outliers), to highlight regions where samples are highly correlated (clusters), or pinpoint populations with a low genetic diversity. A powerful ESDA tool is the open-source software GeoDa (GeoDa, 2005) that allows the exploration and spatial analysis of vector data (Anselin et al., 2006). GeoDa notably offers the possibility to create various maps (quantile, equal intervals, etc.) and to simultaneously analyze attributes with the help of other graphs.

Spatial autocorrelation (i.e. the degree of dependence among observations in a geographic space; SA) is often overlooked in ecological and evolutionary studies despite the fact that many environmental or biological characteristics show spatial dependence among observations, due to intrinsic process of dispersal and mating (Anselin, 1998; Hall and Beissinger, 2014). It is measured by comparing individual values of a defined variable with the mean of that variable in a defined neighborhood. By doing so for each sample, SA measures the degree of values similarities with location similarity. It is thus essential to measure SA in studies involving spatial data, not only because it is a natural phenomenon but also because it violates the assumption of independence required by standard statistical tests, such as student tests or regressions (Legendre, 1993; Wagner and Fortin, 2005). For example, Moran's I, a classic spatial autocorrelation statistic, can be used to estimate the scale/distance of gene flow in the landscape (Hall and Beissinger, 2014). In addition, Local Indicators of Spatial Association (LISA; Anselin, 1995) allows to identify and localize spatial autocorrelation patterns and study the spatial relationship between genetic markers and environmental features (Colli et al., 2014; Stucki, 2014). See Figure 1 for an example. While GeoDa is better to visualize the SA of one variable, it cannot be automated to calculate it for many. For a fast computation of both global and local SA on genetic data, Sambada is handful (Stucki et al., 2016). It can be easily programmed to compute SA on millions of genetic markers and so with different neighborhood sizes and weighting schemes. The decrease of SA with distance can thus be measured using different lags and comparisons can be made between neutral and selected loci. The R package *spdep* can performs similar analyses (Bivand and Piras, 2015).

**Fig 1.**
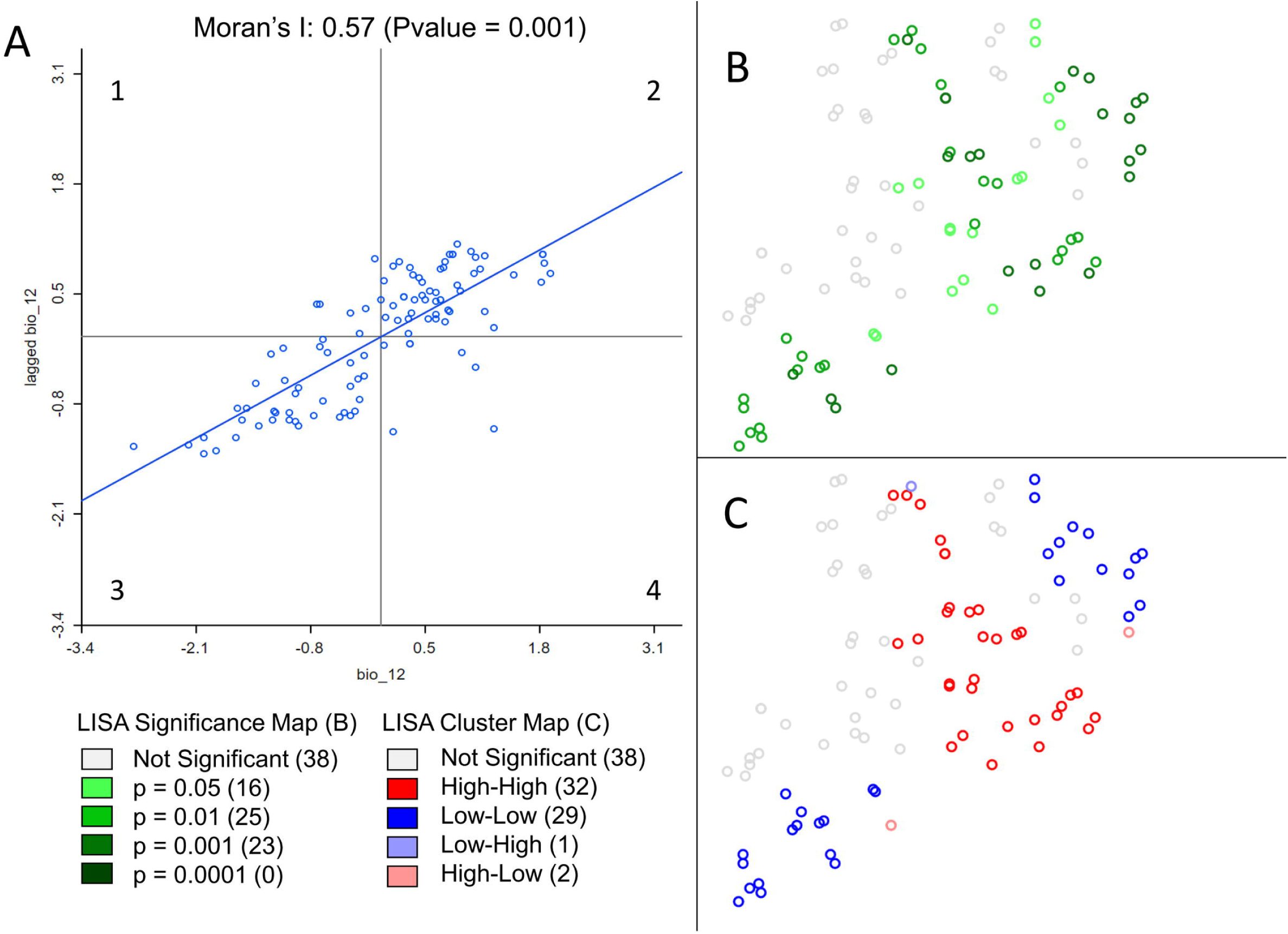
Example of spatial statistic measurement in GeoDa. Results from global and local spatial autocorrelation (SA) were computed on Annual Precipitation at sampling locations of Ugandan cattle (Stucki, 2014). Annual Precipitation was extracted from the WorldClim dataset. In GeoDa, a weight file was created using the 10 nearest neighbors before computing spatial autocorrelation. 999 permutations were performed to assess the significance of both SA measurements. The scatter plot of Global SA (A), measured by the slope of the regression (0.57) displays the standardized precipitation values of each point on the X axis and standardized mean precipitation values of their 10 nearest neighbors on the Y axis. The scatterplot shows a positive correlation between most individuals and their neighbors. In other words, when precipitation is high (low) at a given location, close surrounding locations are more likely to experience high (low) precipitation as well. This positive correlation between neighboring locations is the translation of a clustering of values. On the other hand, significant local SA coefficients (B) are categorized (C) according to the 4 quadrants of the Moran's I plot (A). In contrary to global SA, local SA indicates the location of positive SA or clustering, (High-High - A2, Low-Low - A3), and negative SA or spatial outliers (High-Low - A1, Low-High - A4). Non-significant local SA coefficients are displayed in white.

## Spatial data representation

Maps illustrating the results of an analysis are often more powerful than tables to transmit a result or an idea. However, the creation of efficient maps requires a reflection phase about the graphical representation of the results. Indeed, maps can be too complex to read when too detailed or may be uninformative when too simple. Creating a map first requires choosing an appropriate display type.

Traditional choropleth map, in which the entities are colored according to a scale based on the value of the attribute of interest, can be used in many situations. For example, to represent the membership coefficient of individuals to two populations distributed over a landscape (e.g. as frequently done for population genetic analyses of admixture), one can use a gradient passing through a neutral hue to contrast the two parts of the distribution (i.e. the membership of each individual to one or the other population) (Figure 2). Although most GIS provide colored gradients, it can be useful to understand how to obtain an appropriate color scheme using Color Brewer (Color Brewer 2, 2001). Alternatively, if individuals are grouped into more than two populations, bar charts can be more appropriate. Proportional circles can also be used, for example to indicate absolute numbers of individuals sampled in each population.

**Fig 2.**
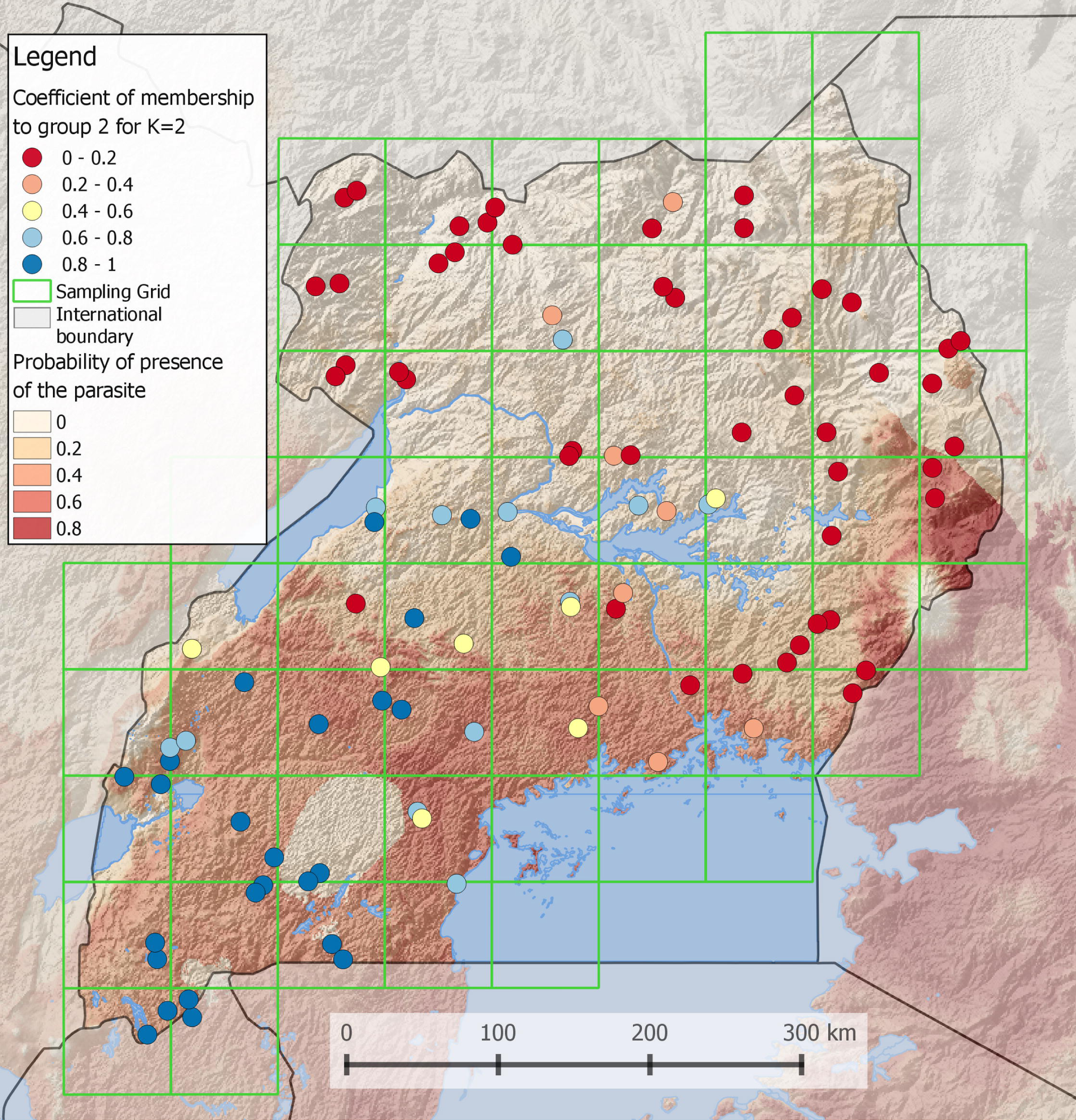
Coefficient of membership to a genetic group of Ugandan Cattles (Stucki 2014). Using the software admixture (Alexander et al., 2009), the most likely number of populations was found to be 2. In this case, it is possible to display the membership coefficient of each individual to one of the two populations. To do so, a gradient obtained from http://colorbrewer2.org is passing through a neutral hue to contrast the two populations. The order of layers in the legend is the same as in the map. In the background, a grid layer of probability of presence of a parasite is shown. A semi-transparent shaded relief is also displayed to reveal the topography. Lakes and international boundaries are overlaid on these raster layers. Ugandan boundaries are highlighted by darkening surrounding countries.

Background layers can then be added to provide more information on the geographic context, such as an aerial image or a DEM to situate the samples. This can potentially be combined with contour lines to compare the elevation from one location to another. One can also highlight the study area by darkening or de-saturating the rest of the map. Regarding points representing individuals or populations, simple shapes should be preferred. Labels should be readable and discarded if not.

Each map must then be edited before being published. Some elements must go along with a map: a legend (to identify the geographical units, or the different statistical classes used) and a scale. In the legend, the message should be simplified by regrouping categories, reducing the number of decimal places and removing unnecessary layers. Furthermore, a frame in a corner of the map, representing the region at a broader geographic scale (zoom out), is useful to situate the study area. Maps should be exported preferentially in .pdf format to keep the vector properties for potential future editions. We provide an example in Figure 2.

Lastly, GIS software are not particularly easy to use when it comes to producing maps iteratively. For example, creating maps of genetic markers under selection used to be feasible manually when the number of genetic markers tested was small. On the contrary, most GEA studies today use hundreds to millions of markers deriving from genomic analyses, and with many of them showing signatures of selection. In such cases, manually producing maps is neither smart nor informative. Computing software such as R should thus be favoured with packages such as *Rgdal* and *Rasters* being very useful and sufficient to import genetic and geodata and to produce basic maps (Bivand et al., 2016; Hijmans and van Etten, 2015).

## Perspective

GIS are powerful tools for molecular ecologists but remain too often underexploited and misused, mainly because of the multitude of GIS software and databases available. We have presented in this paper useful guidelines making it possible for any GIS beginner to appropriate basic functions, to find specific learning resources for biologists, and we proposed a brief state of the art for the use of GIS in biology. However, it is intriguing that, in the big data era, geodatabases are not more frequently used to store and access genetic datasets. They would also speed up queries and reduce disk usage. There are in fact few examples of transformation of NGS data in spatial databases because of the high technicality of such task (Holl and Plum, 2009; Joost and Kalbermatten, 2010; Nandal et al., 2016; Paila et al., 2013; Piry et al., 2016). So far, the most compelling tool is the recently developed open source system TheSNPpit (Groeneveld and Lichtenberg, 2016). It allows for an integration of large genetic datasets in a PostgreSQL environment, which is also the backend of most GIS databases. Interestingly, this tool was mainly developed for breeding programs that already deal with thousands of individuals and millions of SNPs. A game changer that will most likely hit molecular biology in the future.

### Box 1. Sampling design and scale

Sampling design must be carefully chosen depending on the ecological scale of study, i.e. the spatial resolution and the extent of the area under study, and economic constraints. One important way of evaluating sampling strategies is to design them in a GIS environment to guarantee spatial randomness, representativeness or a constrained stratification of the sampling along environmental gradients among others.

Various optimized sampling strategies have been proposed and reviewed in the literature (Balkenhol et al., 2015; Manel et al., 2012; Rellstab et al., 2015; Schwartz et al., 2009). However, and regardless of the design chosen, one must consider sampling density and decide how many individuals will be sampled and then sequenced per location (or population). Indeed, the recent availability of NGS data implies to consider sub-sampling strategies for economic reasons. For example, a sub-sampling procedure using a hierarchical clustering can be applied in order to ensure a regular cover of both environmental and physical spaces (Stucki, 2014). For the former, stratified sampling techniques should be used over a range of climatic variables, previously filtered using a PCA. For the latter, a clustering index is minimized to ensure spatial spread. To ensure the representativeness of the entire area, the sampling can also be achieved using grid cells (see Figure 2). On the other hand, it is important to understand that landscape and population genomic sampling designs are difficult to reconcile (Joost et al., 2013). Indeed, sampling a small number of populations does not necessarily allow estimating changes in frequency along an environmental gradient. Conversely, sampling regularly along an environmental gradient may turn the assignment of individuals to populations more difficult. However, as pointed out by De Mita et al. (2013), for population genetics studies it is preferable to sample a high number of populations with few samples rather than a small number of populations with many samples. In addition, it is better to concentrate the sampling in a smaller area in order to obtain a greater density and higher statistical power (Joost et al., 2010).

Defining a scale of study also raises important questions regarding the relevance of environmental variables used. Indeed, when integrating different datasets (e.g. environmental, topographic, genetic), one must be aware that the spatial resolution of the raster data has to match the sampling density, and this is often not the case. Recently developed satellite imagery or DEMs show a fine resolution and a high accuracy, but the advantage of using high resolution data compared to data at coarser resolution remains under-studied (Cavazzi et al., 2013; Levin, 1992; Marceau and Hay, 1999; Wilson and Gallant, 2000). For example, while intuitively a fine resolution may be ideal, it may hold an excess of details and generate too much noise. Contrastingly, a too coarse resolution only shows generalized properties of the landscape and can have little explanatory power (Cavazzi et al., 2013). On the other hand, when the spatial resolution of the variable is too coarse, nearby samples will retrieve their environmental values from the same pixels (i.e. pseudo-replicates), thus inflating autocorrelation. One solution to this problem is to compute variables at multiple resolutions (Leempoel et al., 2015; Pradervand et al., 2014).

## List of terms

Raster: regular grids of pixels that describe continuous phenomena, retaining information such as color (for aerial images), elevation, temperature.
Vector: points, lines or polygons whose nodes are defined by geographical coordinates and describe discrete phenomenon such as borders, rivers, catchment areas. Vectors are usually stored in Shapefiles (.shp and associated files).
Datum: The datum defines the 3-dimensional sphere used to approximate the earth. It provides a frame of reference to measure coordinates in both geographic and projected coordinate systems.
Geographic Coordinate System: A GCS give the coordinates (i.e. latitude and longitude) of a point as measured from the angles to the center of a defined sphere and meridian.
Projected Coordinate system: A PCS is a projection of the sphere on a flat, two-dimensional surface. Its coordinates (X and Y) are thus consistent and equally spaced.
DEM: Digital Elevation Models are grids of elevation data. Each pixel of that grid is spaced at regular horizontal intervals and contains one value of elevation.
Grain: The grain is the size of a pixel, the smallest unit on a grid. A small grain corresponds to a high spatial resolution.
Extent: The extent is the size of the study area.

